# Fitness and productivity increase with ecotypic diversity among *E. coli* evolved in a simple, constant environment

**DOI:** 10.1101/679969

**Authors:** Dong-Dong Yang, Ashley Alexander, Margie Kinnersley, Emily Cook, Amy Caudy, Adam Rosebrock, Frank Rosenzweig

## Abstract

Community productivity often correlates with diversity. In the microbial world this phenomenon can sometimes be explained by highly-specific metabolic interactions that include cross-feeding and syntrophy. Such interactions help account for the astonishing variety of microbial life, and drive many of the biogeochemical cycles without which life as we know it could not exist. While it is difficult to recapitulate experimentally how these interactions evolved among multiple taxa, we can explore in the laboratory how they arise within one. These experiments provide insight into how different bacterial ecotypes evolve and from these, possibly new ‘species.’ We have previously shown that in a simple, constant environment a single clone of *E. coli* can give rise to a consortium of genetically-and physiologically-differentiated strains, in effect, a set of ecotypes, that coexist by cross-feeding. We marked these different ecotypes and their shared ancestor by integrating fluorescent protein into their genomes. We then used flow cytometry to show that each strain by itself is more fit than the shared ancestor, that pairs of evolved strains are fitter still, and that the entire consortium is fittest of all. We further demonstrate that the rank order of fitness values agrees with estimates of yield, indicating that an experimentally evolved consortium more efficiently converts resources to offspring than its ancestor or any member acting in isolation.

**Importance:** In the microbial world, diversity and productivity of communities and consortia often correlate positively. However, it is challenging to tease apart a consortium whose members have co-evolved, and connect estimates of their fitness and the fitness of their ancestor(s) with estimates of productivity. Such analyses are prerequisite to understanding the evolutionary origins of all biological communities. Here we dissect an *E. coli* consortium that evolved in the laboratory and show that cooperative interactions are favored under continuous glucose limitation because a partnership of ecotypes is better able to scavenge all available resources and more efficiently convert those resources to offspring than any single individual. Such interactions may be a prelude to a special form of syntrophy, and are likely to be key determinants of microbial community structure in nature, including those having clinical significance, such as chronic infections.

## INTRODUCTION

Microbial communities in nature exhibit enormous genetic diversity owing in part to the extreme spatial and temporal heterogeneity of life at the micron scale. This heterogeneity opens up ample opportunities for selection and drift to act differentially on new variants arising by mutation, horizontal gene transfer or arriving via dispersal. Over geologic time-scales these evolutionary forces have enabled microbes to exploit almost every conceivable environment on and in the earth’s crust, where they partition niches according to how they differ in their physiological tolerances and in their electron donor and acceptor preferences (1). Microbes not only partition existing niches, they also create new ones. For example, one microbial taxon may release metabolites that sustain others, who by consuming them relieve product inhibition or render thermodynamically unfavorable reaction sequences favorable (2). Among widely-diverged taxa, interspecies transfer of metabolites can promote stable associations that maximize the amount of energy extracted from the surrounding environment, a phenomenon known as syntrophy (3).

Over ecological time-scales, selection on closely-related members of a bacterial population may result in clonal replacement of successively fitter adaptive lineages (4, 5). Alternatively, selection may result in clonal interference, where adaptive lineages co-exist owing to their similar fitness (6, 7), or in clonal reinforcement (8), where adaptive lineages coexist by virtue of frequency-dependent interactions like differential utilization of public goods (9), mutual inhibition (10), and metabolic cross-feeding (11, 12). These three outcomes are not mutually exclusive: clonal interference could easily occur within each of several stably-coexisting lineages, as could periodic selection of highly-fit novel variants (13). In evolving asexual populations adaptive lineages that coexist owing to frequency-dependent selection are more likely to undergo further diversification than those that are not (14, 15). Over succeeding generations such lineages may become so genetically and physiologically differentiated that they attain the status of ecotypes (16), especially when barriers to horizontal gene transfer arise between them. It is an open question whether such genetically-differentiated ecotypes constitute nascent bacterial species, *sensu* Cohan (17), that in time go on to form stable syntrophic associations.

Helling et al. were among the first to report stable coexistence of multiple genotypes in lab populations of *Escherichia coli* originating from a common ancestor (18). Their finding was especially surprising given that evolution had occurred in a glucose-limited chemostat, a simple, constant environment. Therein, frequency-dependent selection should be less impactful than in serially-transferred batch cultures (19), where populations fluctuate between feast and famine, a regimen known to select for stable polymorphism (20–22), especially in mixed resource (23, 24) and alternating resource environments (25). In the Helling et al. system stable polymorphism was supported by cross-feeding between a primary resource specialist and ecotypes that evolved the capacity to exploit secondary resources (8, 26, 27). The relationship between a primary resource specialist and secondary specialists might not be a simple, one-way, cross-feeding of by-products, but instead a reciprocal interaction based on “resource-service” exchange (28, 29). In one case, the glucose specialist provides overflow acetate for one of the secondary resource specialists, which in turn provides a “service” by scavenging this growth-inhibitory metabolite (8, 26, 27). Less clear are why the numerically predominant primary resource specialist so wastefully metabolizes glucose under carbon limitation, and how widespread this kind of resource-service may be in the microbial world.

While it is widely recognized that microbes secrete myriad compounds, some as overflow metabolites (30–33), some as signaling molecules (34), others as defensive or offensive weapons (35, 36)– all at considerable expense to themselves – it is unclear how often these compounds become growth substrates for other microbes, be they members of the same or different taxa. Moreover, even when they do, there may be evolutionary limits to the stability of cooperative interactions, owing to the risk of becoming dependent on a complementary ecotype and the possibility that cooperative metabolic exchanges reduce overall community productivity relative to a single, autonomous clone (37). To test for these possibilities and to deepen our understanding of conditions favoring evolution of a consortium from a single clone, we re-examined the cross-feeding population described by Helling et al. (18). We evaluated under the original evolutionary conditions, the competitive fitness of each consortium member, the fitness of pairs of strains and the fitness of the consortium as a whole, all relative to fitness of their common ancestor. We then related fitness estimates to estimates of resource consumption and productivity for each ecotype and for combinations thereof. We find that fitness and productivity increase in relation to functional complexity, measured as the number of co-evolved ecotypes present in the system. Thus, just as syntrophic interactions among different taxa can maximize the amount of energy extracted from complex environments in nature, cross-feeding interactions between adaptive lineages arising from a common ancestor can maximize the amount of energy extracted from a simple environment in the laboratory.

## RESULTS

### Co-evolved consortia can be reconstructed in the laboratory

A microbial population originating from a single clone typically becomes polymorphic, with genetically distinct clades sometimes persisting over evolutionary time scales (18, 38–40). These clades have the potential to differentiate physiologically into ecotypes that form consortia supported by metabolic interactions such as cross-feeding (26, 41). To open up a more finely resolved view of how consortia arise under continuous glucose limitation we created GFP-tagged versions of the *E. coli* strains described by Helling et al. (**Table 1**) (18). Flow cytometer data have shown that at steady state a numerically predominant primary resource specialist E3 stably coexists with either, or both, of two secondary resource specialists, E1 and E6 (**Figure 1 A, B**, and **C**). Secondary resource specialists E1 and E6 can also co-exist, but not reproducibly, owing to the low frequency (<1%) of E1 cells in mixed populations at equilibrium (26). At steady state the ratio of E3:E1 was approximately 9:1, whereas that of E3:E6 was approximately 5:1, which accords with previous estimates of relative strain frequency based solely on colony size and antibiotic sensitivity. The evolved strains’ common ancestor, A, was outcompeted by E3 in < 10 generations (**Figure 1 D**).

**Figure 1.**
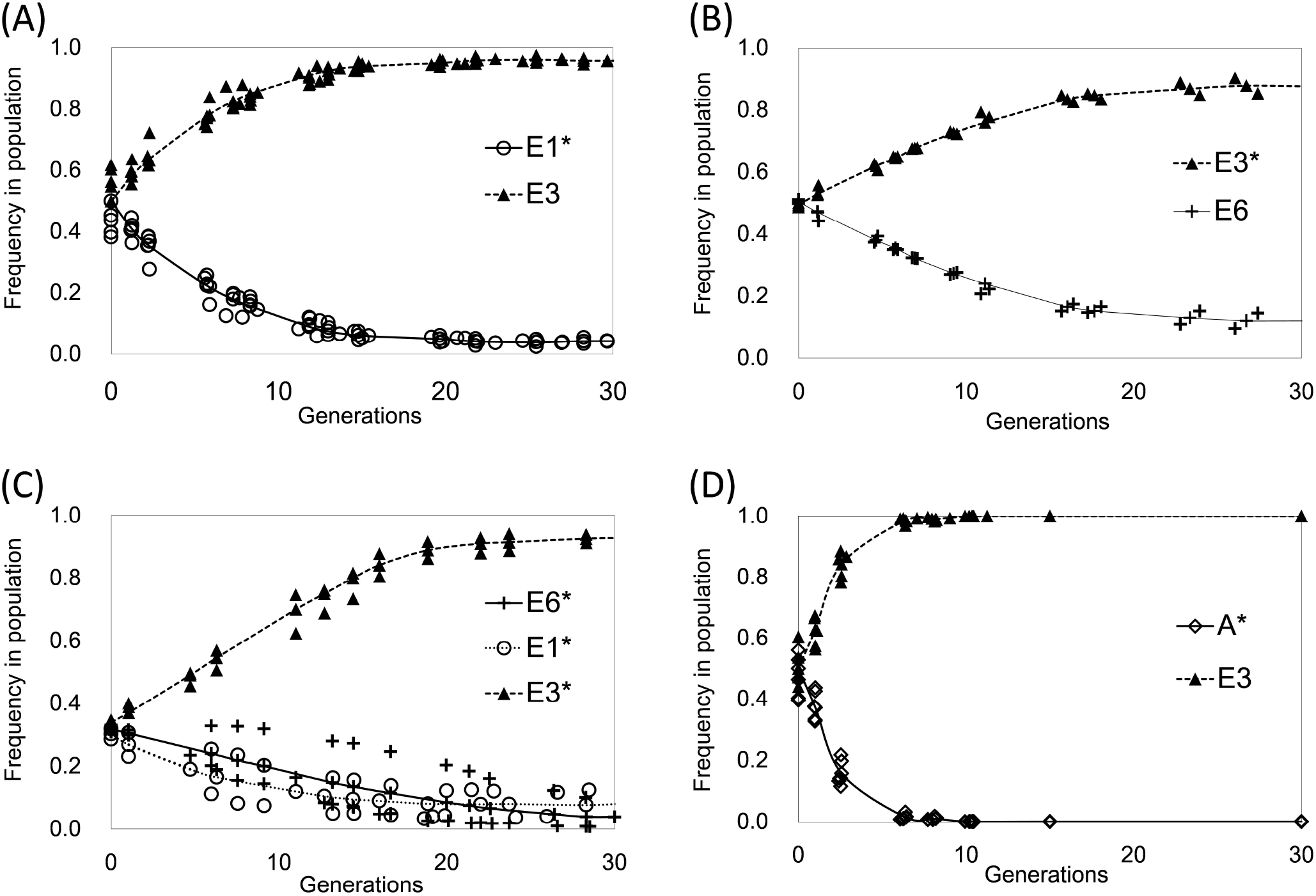
Reconstruction experiments: (A) E3+E1 consortium; (B) E3+E6 consortium; (C) E3+E1+E6; (D) Displacement of the ancestor, A, by E3. Strain frequency in experimental populations was inferred by evaluating the incidence of FP-marked (*) strains by flow cytometry as described in Methods. Individual data points from at least three different experiments are represented.

**Table 1.** Strains and relevant physiological properties

| Strains | Strain Characteristics | Glucose Uptake Rate<sup>1</sup> | Growth on alternative carbon sources |  |  |  |
| --- | --- | --- | --- | --- | --- | --- |
|  |  |  | 0.2% glucose |  | 0.5% acetate |  |
| | | | $\mu_{\max}^2$ | Yield<sup>3</sup> | $\mu_{\max}$ | Yield<sup>3</sup> |
| K12 MG1655 (K) | F- lambda- *ilvG- rfb-50 rph-1* | NA | 0.31±0.00 | 1±0.01 | 0.00±0.00 | 0.04±0.00 |
| JA122 (A) | Derivative of RH 201*, F- *thi1 lacY1 tonA21 supE44 hss1, araD139*, lysogenic for $\lambda$, contains plasmid pBR322Δ5(Amp<sup>R</sup>) | 1.19 ± 0.09 | 0.32±0.01 | 1.15±0.03 | 0.00±0.00 | 0.02±0.00 |
| CV101 (E1) | Derivative of JA122; isolated after 773 generations, Amp<sup>R</sup>, form larger colonies on TA plate. | 1.66 ± 0.06 | 0.38±0.01 | 1.49±0.02 | 0.02±0.00 | 0.08±0.00 |
| CV103 (E3) | Derivative of JA122; isolated after 773 generations, Amp<sup>R</sup>, form small colonies on TA plate. | 2.46 ± 0.16 | 0.26±0.00 | 1.08±0.01 | 0.00±0.00 | 0.00±0.00 |
| CV116 (E6) | Derivative of JA122; isolated after 773 generations, form larger colonies on TA plate, lacks plasmid, Amp<sup>S</sup> | 1.61 ± 0.11 | 0.39±0.00 | 1.27±0.03 | 0.00±0.00 | 0.01±0.00 |

<sup>1</sup> Glucose uptake rate in batch culture ( $\mu\text{mol } \alpha\text{-MG}/\text{min}/\text{gm}$ dry weight biomass) as measured by (Helling et al. 1987). <sup>2</sup>Maximum specific growth rate was expressed in $\text{hr}^{-1}$ . <sup>3</sup>Cell yield was estimated as optical density at $\lambda=550$ nm, and normalized to that of K12 in Davis Minimal medium containing 0.2% glucose. All values are means ± SEM of 3-4 biological replicates.

### The fitness of evolved ecotypes is greater than that of their common ancestor, and the fitness of consortia is greater still

While the transcriptome and the whole genome sequence of every strain in the evolved consortium have been described (8, 27), their fitness and productivity as individuals, and as collectives, have not. To estimate fitness, we first competed the GFP-labeled ancestral strain (A*) against the unlabeled ancestor (A) under evolutionary conditions (0.0125% glucose limitation). We found that the GFP-tagged strain had a slightly negative fitness coefficient (−0.076±0.014). Thus, to compute relative fitnesses we normalized competition coefficients to 1 for A vs. A* to account for the fitness decrement arising from fluorescent protein expression. We competed each evolved ecotype, as well as consortia composed of different ecotypes, against their common ancestor. Every evolved strain was significantly more fit than that ancestor (**Figure 2**) (one-way ANOVA, Tukey’s HSD, *P* < 0.01). Interestingly, evolved strains did not significantly differ from one another in fitness, relative to the common ancestor (one-way ANOVA, Tukey’s HSD, *P* > 0.9), even though they differed markedly from one another with respect to their glucose uptake kinetics, their growth rate and yield on different carbon sources (**Table 1**) (18, 26), as well as in their patterns of substrate utilization established via BIOLOG assay (**Suppl. Materials Figure S1**). Consortia consisting of two evolved ecotypes (E3+E1 and E3+E6) were more fit than their shared ancestor, and E3+E1 was more fit than E1, E3, or E6 monocultures (one-way ANOVA, Tukey’s HSD, *P*<0.05) The three-membered consortium E3+E1+E6 was not only significantly more fit than the ancestor, but also more fit than every evolved strain in monoculture as well as the E3+E6 consortium (one-way ANOVA, Tukey’s HSD, *P*<0.05). Additionally, E3+E1+E6 was more fit than E3+E1 and E3+E6 when compared to them by paired *t*-test (*P*<0.05).

**Figure 2.**
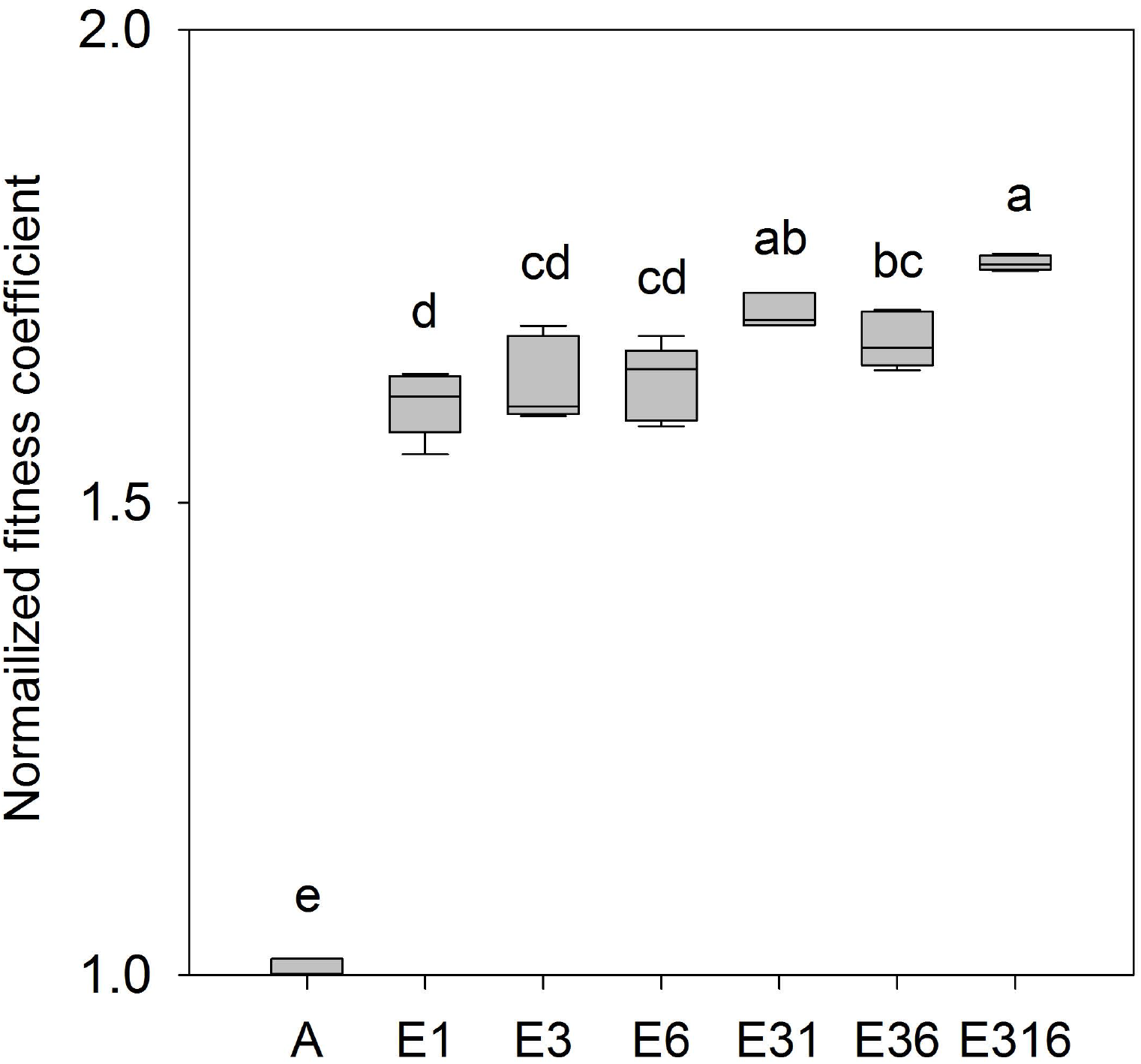
Normalized fitness coefficients of evolved strains relative to their common ancestor: Evolved strains E3, E1, E6 or consortia (E3+6, E3+1, E3+6+1) were competed with GFP-labeled ancestor, A*. Significant differences were identified by one-way ANOVA, and indicated by lower-case letters. Non-identical, lower-case letters indicate significance at P<0.05 (Tukey’s HSD). The boundaries of the box and the whiskers correspond to the 25th and 75th, and the 10th and 90th percentiles, respectively. Median lines are shown in boxes. All fitness estimates were generated from a minimum of four biological replicates.

### Relative fitness differences correlate with differences in productivity

Next, we sought to determine whether the observed fitness differences among strains and consortia mapped onto productivity. Evolved ecotypes, consortia of these ecotypes and their common ancestor were grown to steady state in replicate glucose-limited chemostats and evaluated with respect to yield cell number and dry weight biomass per mL culture. To make our productivity estimates more broadly comparable we also cultured the canonical *E. coli* K12 strain (MG1655) under the same conditions and included its productivity values in our analyses. No significant differences in yield were detected between A and K12 (**Figure 3A, B, C**), whereas each of the evolved strains (E1, E3 and E6) exhibited greater yield cell number than either A or K12 (**Figure 3A**) (*P*<0.05, one-way ANOVA). Surprisingly, when grown as monocultures none of the evolved strains demonstrated higher yield dry weight biomass than their ancestor (**Figure 3B**), indicating that they had evolved smaller cell size, a frequent adaptation by bacteria to chronic nutrient limitation (42, 43). Co-culture of E3+E1 and E3+E6 did result in higher yields than any monoculture (**Figure 3A** and **B**), with differences most pronounced when yield was estimated as dry weight biomass. The median value of biomass in chemostats containing three-members, E3+E1+E6, was highest of all tested and almost two-fold greater than chemostats containing only the common ancestor (**Figure 3B**).

**Figure 3.**
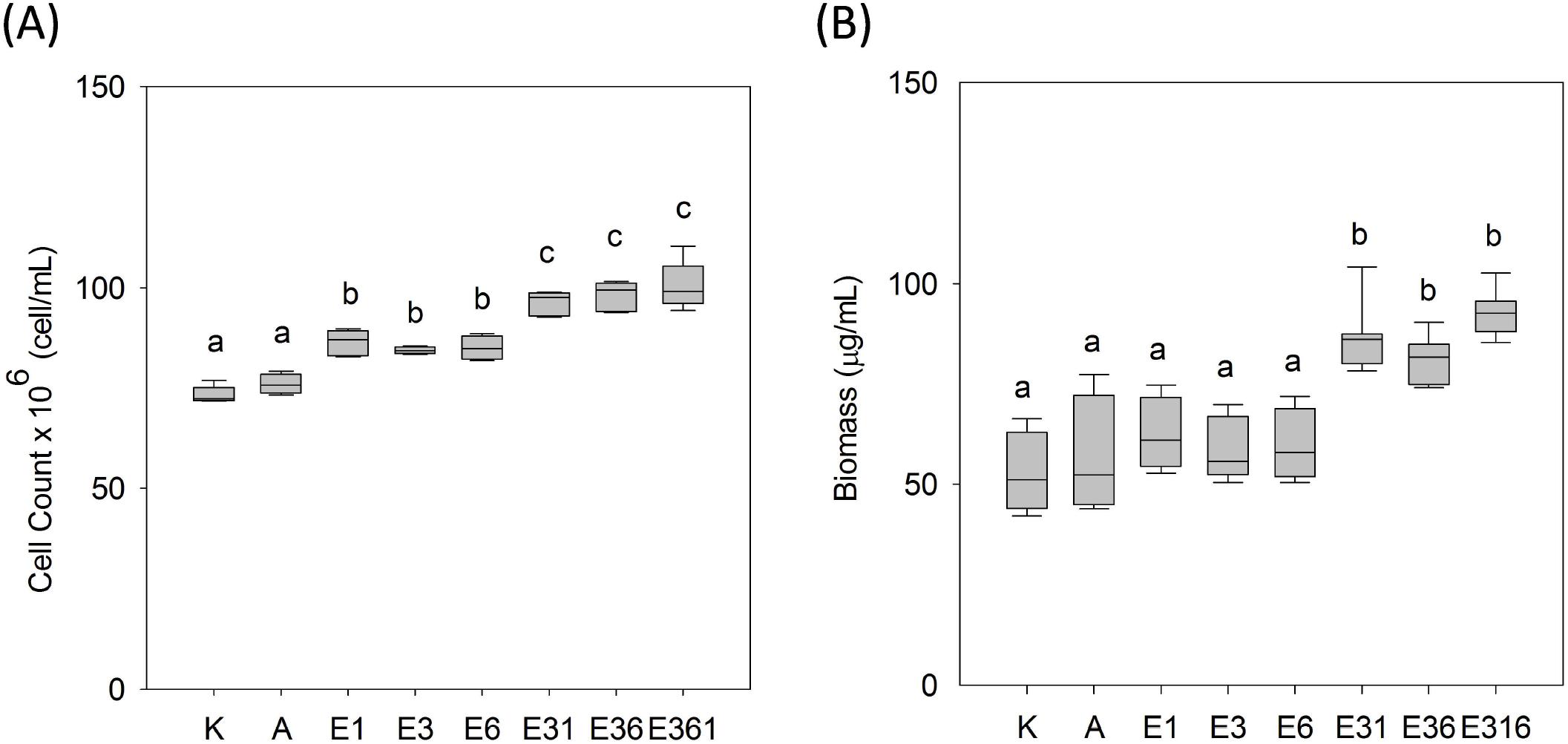
Productivity in steady state chemostats of evolved strains and consortia relative to their common ancestor and to E. coli K12. (A) Yield cells mL-1 (B) yield dry weight biomass in μg mL-1. Significant differences were identified using one-way ANOVA, and are indicated by lower-case letters. Non-identical lower-case letters indicate significance at P<0.05 (Tukey’s HSD). The boundaries of the box and the whiskers correspond to the 25th and 75th, and the 10th and 90th percentiles, respectively. Median lines were shown in boxes. All productivity estimates were generated from a minimum of three biological replicates.

### Residual metabolite levels differ between monocultures and consortia

When grown to steady state under continuous glucose limitation, the ancestral strain and the evolved ecotypes significantly differed with respect to residual metabolite concentrations, with differences previously noted among monocultures (26) for residual glucose and acetate largely confirmed here (**Table 2**). Chemostat populations of K12 or A left the highest concentrations of residual glucose, while those of evolved strain E3 left the lowest, less than 10% of the ancestral strain’s mean value. This finding is consistent with E3’s exceptional capacity to scavenge limiting glucose (**Table 1**). A one-way ANOVA did not uncover significant differences in residual glucose levels among monocultures or consortia of the evolved ecotypes. However, direct comparisons using paired *t*-tests showed that residual glucose differed between E3 and E1, and between E3 and E6 (*P*<0.05). E1 left no residual acetate, confirming that it is an acetate scavenger (26, 44). No residual acetate was found in consortia containing the E1 ecotype (E3+1, E3+1+6), indicating that E1 completely consumes this secondary resource.

**Table 2.**
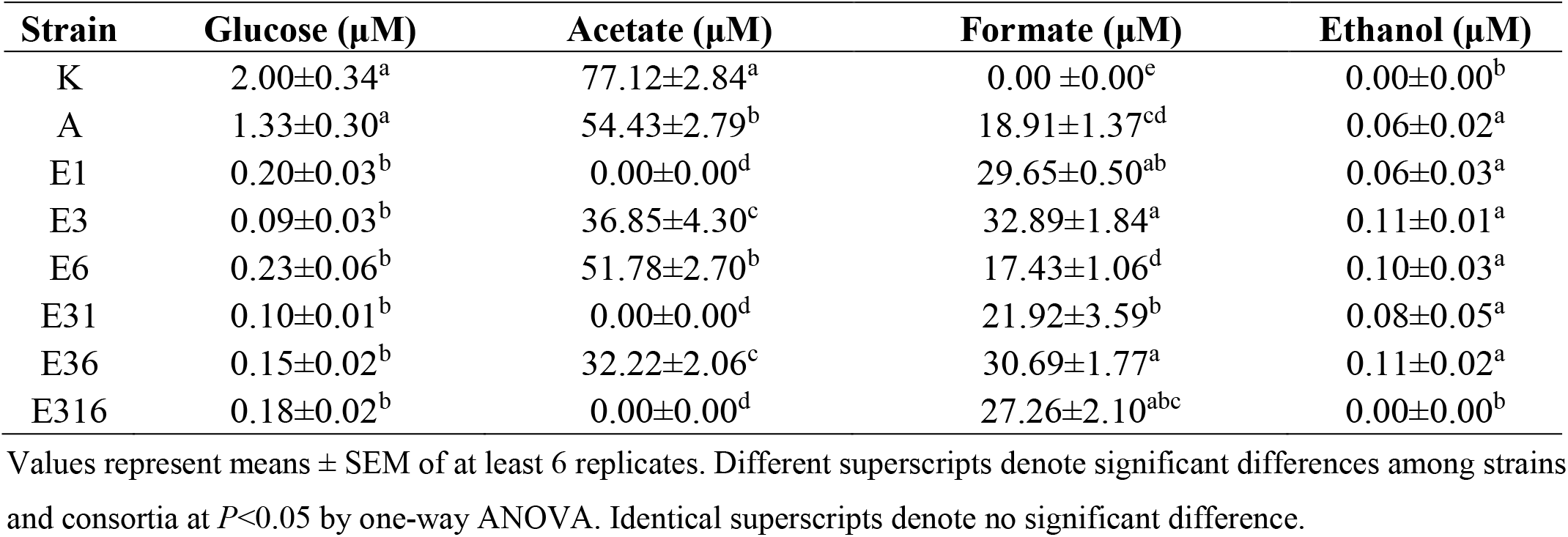
Residual steady-state metabolite concentrations

We also investigated whether other residual metabolites might be implicated in cross-feeding. Unlike the canonical K12 strain, formate overflow was observed in all single- and multi-strain chemostat cultures (**Table 2**). Residual formate levels in E6 monocultures were comparable to those in A, but significantly less than those in chemostat cultures of the other evolved ecotypes as well as the consortia (one-way ANOVA, Tukey’s HSD, *P*<0.05). Residual ethanol was detected in all but K12 monocultures and cultures of the three-membered consortium (**Table 2**). 2-hydroxyglutarate, the reduced product of TCA intermediate 2-oxoglutarate, was several-fold higher in ancestral strain than in K12 or any evolved strain or consortium; similarly, TCA intermediate, *cis*-aconitate, was several-fold higher in chemostats populated by A than in those populated by K12 or any of the other evolved ecotypes. Five unknown metabolites were also identified by LC-MS (**Table S3**), and most of these showed lower residual values in consortia than in monocultures, especially those of K12 and A.

### BIOLOG assays reveals additional ways in which the experimentally evolved ecotypes are phenotypically differentiated

To determine whether the evolved ecotypes differed in other ways that open up the possibility for metabolic interactions we tested their substrate utilization patterns using the BIOLOG platform: a multiplexed phenotyping system carried out in 96-well format on batch cultured cells. Relative to the ancestor and to the other evolved strains, acetate specialist E1 grew best on acetate, pyruvate, malate and fumarate (**Figure S4A**), indicating a greater capacity for TCA cycle metabolism as well as for aerobic and anaerobic respiration (**Figure S5**). We also discovered many instances where evolved ecotypes differentially assimilated carbon, nitrogen, phosphorous and sulfur growth substrates (**Figure S4 A-D**). For example, relative to E1 and E3, ecotype E6 grew best on diffusible intermediates such as 6-phosphogluconate, D-2-phosphoglyceric acid, β-glycerol phosphate, glycerol and D,L-a-glycerol phosphate (**Figure S4C, D**), indicating that this ecotype has greater assimilative capacity along the pentose phosphate and Entner-Doudoroff pathways (**Figure S5**). The existence of so many complementary metabolic capacities, coupled with additional evidence that E6 is especially adept at using multiple phosphorylated 3-carbon intermediates, opens up possibilities for cross-feeding interactions not easily inferred by direct assay of residual substrate concentrations in spent chemostat media.

## DISCUSSION

### Ecological complexity in the microbial world stems from chemical interactions

Microbes engage in every conceivable form of biotic interaction, ranging from intraspecific and interspecific competition (45, 46) to predation (47, 48), to a variety of cooperative interactions that encompass communal feeding (49, 50), detoxification of growth-inhibitory compounds (51), and cross-feeding (52, 53). Played out against a backdrop of spatiotemporal variation in abiotic factors such as pH, redox potential, pO_2_, inorganic ions, light, and temperature (54–61), biotic interactions give rise to the structure and dynamics of microbial communities in nature. Of those interactions that are cooperative, all are ultimately mediated by chemicals secreted into the extracellular environment, be they quorum-sensing molecules (62), degradative enzymes (63) and polysaccharides (64), or overflow metabolites (65), including hydrogen (66).

Synthetic biology has proved to be a powerful tool for unravelling microbial interactions driven by secreted exometabolites (67, 68), leading for calls to standardize fabricated ecosystems in order to systematically dissect signaling and metabolic exchange in natural communities (69). For example, cross-feeding can be genetically engineered, leading to the discovery that in *E. coli* compartmentalization of glucose catabolism and overflow metabolite catabolism in different cellular compartments enhances productivity, whether in batch, chemostat or biofilm culture (70). Similarly, the distribution of two anabolic pathways between a genetically engineered yeast co-culture with an engineered *E. coli* mutualist resulted in enhanced natural product formation (71). Multiple groups have created complementary amino acid auxotrophs and locked them into obligate mutualisms (72–75). These systems have been used to evaluate the fitness costs and stability of this type of microbial interaction, as well as the genetic changes that ensue after it is established (76, 77).

Nothwithstanding insights offered via synthetic biology, few studies have reported on how cross-feeding spontaneously emerges in the laboratory, especially under constant resource-limiting conditions. Models based on such studies open up the possibility for investigating the genetic and ecological conditions in nature that favor cross-feeding (44), a phenomenon that occurs in both clinical (78, 79) and industrial settings (80–82), with important consequences for therapy and bioproduction. The spontaneous evolution of cross-feeding within one taxon may also be a prelude to syntrophy, which typically involves unrelated taxa, and has most often been documented in environments where electron acceptors such as oxygen, sulfate or nitrate are scarce, and complex electron donors are first fermented by one taxon to educts like H_2_, formate or acetate, then further oxidized by partner taxa (28, 83–87).

Theory suggests syntrophy may be rare, owing to inherent risks in obligate metabolic codependency as well as to diminished efficiency of substrate use by a collection of strains relative to one that is autonomous (37, 88). Nevertheless, in microbial populations originating from a single clone, the appearance of genetic diversity can sometimes be ascribed to the evolution of different lineages pursuing complementary metabolic strategies (18, 26, 38, 39, 89). Furthermore, when such complementary strategies have been engineered, rather than evolved via natural selection, the resulting synthetic consortia exhibit increased productivity (70), stability and/or fitness (74). Taken together, these findings suggest that metabolic interactions in the form of cross-feeding, whether within or between species, likely play a key role in determining microbial community structure in nature (90).

### Ecotypic differentiation and partnership in a simple constant environment

When grown in isolation the ancestral and co-evolved strains in these experiments differ with respect to their growth rate and yield in batch culture, their glucose uptake kinetics, their relative susceptibility to acetate inhibition, and their propensity to release or to take up a variety of overflow metabolites (18, 26, 44). Because these strain-specific physiological differences are associated with genetic differences across scores of loci (8), E1, E3 and E6 should be regarded as ecotypes that have undergone adaptive diversification (91) and that may have the potential to evolve further into bona fide species (17, 92).

All things being equal, when a group of organisms competes for a single limiting resource, the variant best able to scavenge that resource to the lowest concentration should prevail (93–95). Interestingly, under glucose-limitation ecotype E3 was not more fit than the other evolved ecotypes, even though E3 exhibited superior glucose uptake kinetics and consistently rose to highest frequency in co-culture (26). E3 was originally identified as a small colony variant in the evolving population, and later shown to be avid for limiting glucose, but wasteful in its use, releasing overflow metabolites that construct new niches for other variants to exploit (26). Ecotypic variants E1 and E6 exploit niches created by E3, and the fitness of E3+E1, E3+E6 and E3+E1+E6 consortia all exceed fitness of the ancestor. Median fitness values for partnerships were greater than those of all singletons, ancestral or evolved, and the fitness of E3+E1 and E3+1+E6 consortia were statistically greater. That E3+E6 was not significantly more fit than E3 or E6 in monoculture can be attributed to the fact that E6, unlike E1, does not consume an inhibitory overflow metabolite (acetate) but rather exhibits high affinity for 3-carbon compounds like glycerol-3 phosphate. Yield cell number in consortia exceeded that of the evolved ecotypes, their common ancestor and *E. coli* K12. This finding is consistent with theory indicating that there is a second way for an ecotype to prevail under simple nutrient-limiting conditions: namely, to become efficient at converting the limiting resource to offspring (96, 97). In our model, consortia are evolutionarily advantaged because the collective excels at both strategies: scavenging the limiting resource to low concentrations and efficiently converting that resource to offspring.

Ecotypes E3 and E1 exhibit a reciprocal interaction that takes the form of “resource-service exchange” (29, 98). E3 scavenges glucose to residual levels that are inaccessible to E1 but excretes a “resource,” overflow carbon in the form of acetate, to which E1 has preferential access. The E1 ecotype in turn provides a “service” to ecotype E3, by consuming this growth-inhibitory metabolite (99) to which E3 is especially sensitive (44). Under glucose limitation the fitness of E3+E1 consortia exceeds that of ecotypes E1 and E3, validating the conclusion that their coexistence is a form of mutualism that we have previously termed “clonal reinforcement” (8).

The role played by ecotype E6 is more subtle: E6 exhibits superior growth on 3-carbon metabolites and amendment of glucose-limited E3+E6 consortia with glycerol renders E6 numerically dominant over E3 (26). However, a resource-service interaction between E3 and E6 based on overflow glycerol or glycerol-3 phosphate may be difficult to achieve, as neither of these compounds are growth-inhibitory. As relative fitness of these two ecotypes does not significantly differ, they may coexist via clonal interference, albeit stabilized by E3 ➔ E6 cross-feeding of 3-carbon exometabolites – a possibility we aim to test in future experiments. Whatever the mechanism, populations having the highest overall mean fitness were those that were most genetically complex, an observation consistent with a large body of literature indicating positive correlations between community and ecosystem diversity, stability and productivity (100–103).

### Could cross-feeding within a population of facultative anaerobes lead to syntrophy?

Syntrophy has been described as a strategy whereby bacteria use coordinated metabolism to extract the maximum amount of energy from scarce resources under anoxic conditions (28, 84, 86). Under these conditions, different (syntrophic) steps carried out in different cellular compartments make thermodynamically unfavorable reactions favorable. Here we see a facultative anaerobe that can mineralize glucose (18) and genetically diversify into sub-populations that compartmentalize this process. The cross-feeding interactions established thereby are based one clone behaving like an anaerobe, even in the presence of oxygen. Why be avid but wasteful when resources are limited, and why does this collective outcompete an autonomous generalist?

Multiple hypotheses have been advanced to explain why bacteria and other cells achieve maximum growth rate by switching from respiration to fermentation (104). One idea is that complete oxidation of glucose via the TCA cycle results in overproduction of reduced co-factors, especially NADPH, and that the switch enables cells to adjust redox status (105, 106). Other hypotheses include economic arguments based on the cost of synthesizing TCA enzymes (107), electron transport proteins (108) and/or the possibility that (inner) membranes may become space-limited for the integration of electron transport proteins (109). In any case, fermentation allows for faster ATP production per unit membrane area (104).

An adaptive phenotype shared by all consortium members is the upregulation of outer and inner membrane proteins involved in glucose scavenging, notably LamB and the MglBAC complex (27). These membrane proteins are most dramatically overexpressed in ‘avid but wasteful’ E3 (27), placing a premium on membrane ‘real estate’ that could otherwise be dedicated to electron transport proteins. E3 harbors unique *de novo* mutations in lipoamide dehydrogenase (*lpd*) which plausibly reinforces its fermentative strategy (8). And in fact, under aerobic conditions E3’s redox state is like that of an oxygen-limited cell (44), a state wherein Enzyme-Flux Cost Minimization models predict that yield-inefficient pathways like acetogenic fermentation can provide a substantial growth advantage (97). Our conceptual model (**Figure 4**) is one in which E3’s glucose-avid phenotype is achieved at the cost of releasing overflow metabolites that E3 cannot access, which creates secondary niches for ecotypes that can. Coordinated metabolism via cross-feeding then helps the consortium as a whole to maximize ATP production and yield, while simultaneously minimizing enzyme and intermediate concentrations (107, 110).

**Figure 4.**
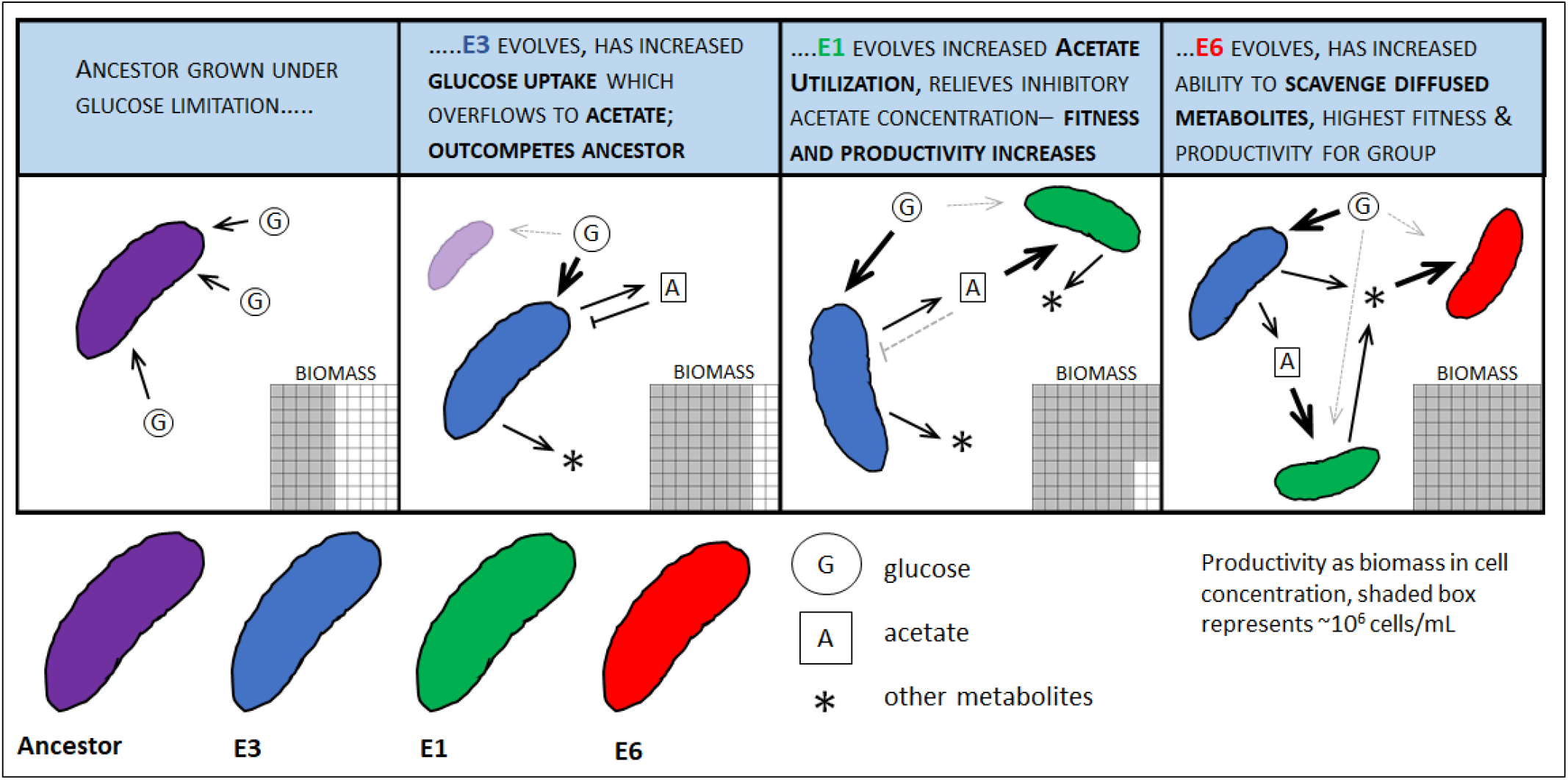
Conceptual model for the spontaneous emergence and maintenance of a cross-feeding consortium from an initially clonal population

### Cross feeding interactions are likely ubiquitous

Syntrophic cross-feeding interactions exist not only between anaerobes, but also between anaerobes and aerobes (111), plants and microbes (112), and between animals and their gut microbiota (113, 114). Cross-feeding may even play a role in chronic microbial infection (115), as complementary auxotrophies have been observed during early phase of *P. aeruginosa* cystic fibrosis lung infections (116, 117). Whether polymorphic microbial populations in these ‘natural experiments’ are collectively more fit and/or more productive than their ancestors, or one another in isolation, has yet to be determined. Crossfeeding is a conspicuous feature of organisms that have differentiated multicellularity, with different nutritive functions segregated into different compartments. And it also arises within genetically heterogenous tumors (118–120) as well as between tumor cells and the surrounding normal tissue to which they are related (121, 122). Only now are researchers beginning to elucidate the genetics and demography of how these metabolic interactions arise and persist in cancer and in chronic infections, which may ultimately help to explain their resilience and their resistance to drug therapy (123). Lastly, the observation that diverse microbial communities, whether engineered or arising spontaneously, exhibit higher fitness (74) and productivity (70) than those that are less so opens up the possibility of engineering novel communities that capitalize on crossfeeding to achieve complex and/or energetically difficult tasks in a stable, robust way (124–127).

## MATERIALS AND METHODS

### Strains, media, and culture conditions

All *E. coli* strains (**Table 1**), including those <u>F</u>luorescent <u>P</u>rotein (FP)-tagged strains used in flow cytometry experiments were archived as 20% glycerol stocks at −80°C. For brevity, ancestral strain JA122 is hereafter designated as A, and its evolved descendants CV101, CV103, and CV116 are designated E1, E3, and E6, respectively. Davis Minimal Medium, DMM (7 gL^−1^ K_2_HPO_4_, 3 gL^−1^ KH_2_PO_4_, 1 gL^−1^ (NH_4_)_2_SO_4_, 0.1 gL^−1^, MgSO_4_·7H_2_O, 0.001 gL^−1^ thiamine-hydrochloride) was used for all chemostat cultures with 0.025% glucose added for batch cultures and 0.0125% glucose for chemostats (128). Maximum specific growth rates (μ_max_) presented in Table 1 were assayed at 30 °C using a Synergy HTX multi-mode microplate reader (BioTek, Winooski, VT) using 48-well plates, each of which contained 0.5 mL Davis Minimal medium amended with 0.025% glucose (w/v). μ_max_ was calculated by following the change in Absorbance at λ=550 nm over time and calculated as previously described (18).

Chemostat inocula were prepared from single colonies on Tryptone Agar (TA; 10 gL^−1^ tryptone, 5 gL^−1^ NaCl, 14 gL^−1^ agar, with or without ampicillin) that were grown overnight at 30 °C in liquid batch culture. Continuous culture was carried out using the Infors Sixfors Bioreactor System (Infors AG CH-4103 Bottmingen/Switzerland). Chemostats were aerated with sterile ambient air, and maintained at 30 °C at a dilution rate, D ≈ 0.2 hr^−1^ for approximately 70 hours (~15 culture generations). Cultures were sampled at prescribed intervals and assayed with respect to cell density by measuring absorbance at 550 nm, and by counting Colony Forming Units (CFU) following serial dilution and plating on Tryptone Agar. Colonies were further scored with respect to colony size (<u>L</u>arge/<u>S</u>mall) and to antibiotic sensitivity (<u>R</u>esistant/<u>S</u>ensitive).

### GFP labeling of E. coli strains

To estimate the relative frequency of different *E. coli* strains as they were competed in chemostats we followed the change in frequency of Green Fluorescent Protein (GFP)-tagged strains over successive generations. An expression cassette containing GFP under the constitutive PA1 promoter was integrated into the genomes of the ancestral and evolved strains using a modified Tn7 delivery system (129). Details of pGRG36-Kan-GFP plasmid construction are described in **Supplementary Materials S1** and primers used in **Supplemental Table S2**. Transposition of the GFP cassette to the attachment site (*attTn7*) at the 3’ end of *glmS* was confirmed by sequencing and epifluorescence microscopy. Our novel plasmid pGRG36-Kan-GFP, which enables rapid GFP-tagging of either the *E. coli, Salmonella* or *Shigella* chromosomes, is available via Addgene (#79088, Cambridge MA, USA)

### Reconstruction experiments

Earlier work had shown that the evolved consortium could be reconstructed as various combinations of the original strains cryopreserved as −80°C glycerol stocks (26). Three groups of reconstruction experiments were carried out under previously described conditions (26), except that in each case one of the strains was GFP-labeled (denoted by asterisk *) to allow monitoring of population frequency by flow cytometry. Group A reconstruction experiments used E3 and GFP-labeled E1*. Group B reconstruction employed E6 and GFP-labeled E3*. In the three-strain reconstruction experiment C, one of the three evolved strains (E1, E3 or E6) was alternatively labeled in each reconstruction. Three to six biological replicates were performed, and the strains’ relative frequencies were estimated by flow cytometry.

### Fitness estimates

Relative fitness of each member of the evolved community, as well as various combinations of those members, was estimated under evolutionary conditions by competing them against the GFP-tagged version of their common ancestor, JA122 (denoted as A*). For each experiment, single colonies of A* and the strain(s) to be analyzed were suspended in 3 mL Davis Medium (0.025% glucose) and cultured overnight at 30 °C, shaking at 150 rpm. Overnight cultures were inoculated into individual SixFors bioreactors containing 300 mL Davis Medium (0.0125% glucose). Reactors were initially run as monocultures, operated at 30 °C in batch mode for 8-10 hours until the cell population attained an absorbance at 550_nm_ (A_550_) of 0.1, whereupon they were shifted to continuous mode at dilution rate, D≈0.2 hr^−1^, and grown to steady-state, approximately three volume changes.

To initiate competition between two strains, the labeled ancestor, A* and one of its descendants: E1, E3 or E6, equivalent cell numbers from each steady-state monoculture were mixed in a 300 mL working volume bioreactor to produce a starting A_550_ of ≈ 0.1, the typical density achieved by strains at steady-state under the limiting glucose concentrations employed by Helling et al. 1987 (18). Once combined, which established time-zero (T_0_), the reactor was placed under continuous glucose limitation and run at D ≈ 0.2 hr^−1^ for 15-20 generations. To initiate competitions between the labeled ancestor, A* and one of several possible multi-strain consortia, monocultures of each evolved strain were first grown as described above, then mixed to reconstitute the consortium at previously described steady-state ratios (18, 26): E3+E1=5:1; E3+E6=4:1; E3+E6+E1=5:1:1. Each consortium was cultured under continuous glucose limitation for 12 hours (~ two volume changes) before adding an equal number of the GFP-labeled ancestor, A* to produce a T_0_ starting density of A_550_ ≈ 0.1. All competition experiments were performed under conditions identical to those under which the consortium had evolved. In both sets of experiments strain frequency dynamics were followed by flow cytometry of chemostat samples that had been taken every 4-5 hours, mixed with 0.5 mL DMM containing 50% glycerol (v/v), and stored at −20 °C until analysis.

### Flow cytometry

To estimate relative abundance of GFP-labeled and unlabeled cells, chemostat samples were diluted 50-fold in 1% (w/v) NaCl and analyzed on an Attune NxT Acoustic Focusing Flow Cytometer (Thermo Fisher, Waltham MA, USA), with a minimum of 10,000 cell events per collected sample. A 50 mW 488 nm laser was used to reveal scatter and fluorescence signals. The Forward Scatter (FSC) signal was detected via a photo-diode detector, while the Side Scatter (SSC) and fluorescence signals were detected by Photo Multiplier Tube detectors. GFP was detected on a 530/30 bandpass filter refined with a 503 nm long pass dichroic filter. During instrument setup, the threshold was set in SSC, which is preferred for bacterial detection, rather than the typical FSC. Bacteria were first gated in the FSC/SSC dot plot to include the bulk of the cells, while excluding debris and large clumps. Single cells were gated and viewed in either a histogram or dot plot of fluorescence vs SSC. A gate was placed around the positive signal and the resulting percentage recorded. For controls, pure cultures of GFP-labeled and un-labeled *E. coli* JA122 were run to view the background fluorescence. The per-generation fitness coefficient was calculated as the slope of the linear regression ln(experimental/reference) as a function of elapsed generations; cell generations elapsed equals (time x dilution rate)/ln2 (ref. 130). Values were calculated as normalized means ± standard deviations from at least four independent experiments. To verify the presence of each experimental strain and to check for contamination during the competition, culture samples were also plated on TA (with or without ampicillin) and scored for Amp^R^/Amp^S^, and Large/Small colony size as previously described (18).

### Cell count and biomass dry weight

Yield parameters were estimated on individual strains and consortia grown to steady state (three-to-five volume changes) in 0.0125% (g/v) glucose-limited chemostats. Cell density per mL was estimated by diluting chemostat cultures 50-fold with 1% (w/v) NaCl then subjecting the resulting suspension to flow cytometry, as described above. The per mL cell number was estimated by the events per μL recorded in the gated region where debris and large clumps were excluded. Dry weight biomass was estimated by filtering a 200 mL aliquot of cells onto tared 47 mm diameter HNWP membranes (Hydrophilic Nylon White Plain 0.2 μm pore size, Millipore, Burlington MA, USA), drying the filters at 65 °C for 24 hours, then re-weighing them using a semi-micro analytical balance (XPE26C, 1 μg readability, Mettler Toledo, Columbus OH, USA).

### Residual metabolites

For each strain and consortium, 50 mL of medium was sampled at steady state from three to four independent chemostat runs. Each sample was filter-sterilized first through a 0.2 μm HNWP membrane (Millipore) filter, then through a 0.22 μm cellulose nitrate syringe filter. Filtrates were stored in sterile 50 mL Falcon tubes at −80°C prior to analysis. Samples for liquid chromatography-mass spectrometry (LC-MS) analysis were re-filtered using a 0.2 μm PES sterile syringe filter and stored at −20°C.

#### Residual glucose

Before assaying glucose, which is at a low concentration in steady-state chemostats, 10 mL of sterile filtrate were concentrated 20-fold by lyophilization, then resuspended in 0.5 mL sterile Millipore water. Extracellular glucose was assayed enzymatically using the D-Glucose Kit (#10716251035, Roche, Basel, Switzerland), based on the production of NADPH, which was assayed spectrophotometrically at 340 nm.

#### Residual ethanol, acetic acid and formic acid

Concentrations of acetic acid and formic acid in culture supernatants were analyzed using an Agilent (Santa Clara CA, USA) 1100 HPLC system equipped with a 7.7 × 300 mm Agilent Hi-Plex H column with 8 μm packing equipped with a 5 mm × 3 mm guard cartridge. A 20 μL volume of sample was injected, using 10 mM sulfuric acid as running buffer. The column compartment was held at 65 °C and the flow rate was 0.6 mL/min. Analytes were detected using a diode array detector (DAD) set to 210 nm. Concentrations were determined by comparison to spiked-in standards. Residual ethanol concentration was determined enzymatically, as previously described (131).

#### Other organic acids

Cell supernatants were analyzed using acidic reverse phase LC-MS on an Agilent Technologies 6540 Q-TOF as previously described (132).

### BIOLOG phenotyping

Utilization of 190 carbon sources, 95 nitrogen sources, 59 phosphorous sources, 35 sulfur sources and 95 nutritional supplements was conducted by Biolog Inc. Phenotype Microarray Services (Hayward CA, USA) using the OmniLog system V. 1.5 (133). Eight 96-well plates, each containing 95 substrates and one negative well (PM1-8) were inoculated with either the A, E1, E3 or E6 strain and assayed in duplicate over 48 hours at 30°C. Substrates for which the absolute value of the background-corrected average well height for test (E1, E3 or E6) minus the average well height for reference (A) strains for both replicates passed a static threshold determined by the company were considered differentially utilized. Positive differences were scored as gained phenotypes and negative differences were considered phenotypic losses. Clustering of average well height values was done using the MeV TM4 software suite (available at https://sourceforge.net/projects/mev-tm4/).

## ACKNOWLEDGMENTS

We are grateful to Karen Schmidt and Pam Shaw for technical assistance. We thank Matt Herron, Eugene Kroll, Pedram Samani and Gavin Sherlock for fruitful discussion and for editorial comments on the manuscript. DDY, MK, AA and FR were funded by NNX12AD87G-EXO from NASA; FR was additionally funded by NASA grants NNH13ZDA001N-EXO and NNH13ZDA017C, the Georgia Tech node of the Astrobiology Institute. AC and AR were funded by the Canadian Institutes for Health Research, the National Science and Engineering Research Council of Canada, the Canadian Foundation for Innovation and the Leaders Opportunity Fund. AC is the Canada Research Chair in Metabolomics for Enzyme Discovery. Collection and processing of flow cytometry data were supported by an Institutional Development Award (IDeA) from the National Institute of General Medical Sciences of the National Institutes of Health under grant number P30GM103338, as well as by funding from the MJ Murdock Charitable Trust.

## COMPETING INTERESTS

The authors declare that they have no conflict of interest.

